# Complex Social Vocalizations are Integrated into the Bat Wingbeat Cycle during Flight

**DOI:** 10.64898/2026.02.12.705650

**Authors:** Simon F. Bauerle, Leonie S. Reetz, Annette Denzinger

## Abstract

Acoustic communication plays a central role in animal social behavior and exhibits remarkable complexity in various taxa. In bats, the emission of echolocation calls is tightly coupled to wingbeat and respiratory rhythms. However, little is known about how complex social vocalizations are coordinated with the mechanics for powered flight. We investigated how complex social calls are integrated into flight of wild Nathusius’ pipistrelles (*Pipistrellus nathusii*). Using synchronized audio and video recordings, we quantified wingbeat rhythms during free flight and incorporated these data into a kinematic inference model to reconstruct the timing of social calls relative to the wingbeat. We found that echolocation call emission is coupled to the upstroke phase of the wingbeat. Moreover, social calls span three wingbeat cycles during flight and are embedded within a continuous sequence of echolocation calls. While motif durations scale with syllable number, motif onsets are phase-locked to narrow, preferential phases of the wingbeat cycle. Our findings provide evidence that complex social calls in bats are integrated into the wingbeat during flight, demonstrating that vocal-locomotor coupling extends beyond echolocation to shape the structure of social communication. By mitigating the high energetic costs of vocalizing, biomechanical integration might have facilitated the evolution of complex vocal communication in bats.

## Introduction

Acoustic communication is widespread across the animal kingdom, underpinning key social behaviors, such as mating and territory defense (Bradbury & Vehrencamp, 2011; Catchpole & Slater, 2008; Gillam & Fenton, 2016). In select vertebrates, most notably songbirds and whales, this acoustic communication has evolved to form an elaborate repertoire of complex vocalizations. This vocal refinement has been critical for providing the communicative basis necessary to facilitate a broad diversity of sophisticated social interactions (Smotherman et al., 2016).

Bats constitute the second-largest mammalian order and differ from all other mammals by their capacity for powered flight, and with the exception of Old World fruit bats, by the use of a laryngeal echolocation system (Altringham, 2011; Denzinger & Schnitzler, 2013; Fenton & Simmons, 2014; Gillam & Fenton, 2016; Kunz & Fenton, 2005). Despite being highly vocal in terms of echolocation, their capacity for highly complex social communication was long overlooked (Smotherman et al., 2016).

Historically, research focused almost exclusively on echolocation calls which are used for spatial orientation in flight and prey capture. However, a growing number of studies have elucidated a rich diversity of bat social behaviors, revealing that many species possess elaborate vocal repertoires (Barlow & Jones, 1997; Behr et al., 2004; Jahelková et al., 2008; Knörnschild et al., 2017; Schmidbauer & Denzinger, 2019). Notably, some of these vocalizations exhibit structural complexity comparable to avian song (Smotherman et al., 2016), and are even maintained through vocal learning (Fernandez et al., 2021; Knörnschild, 2014; Knörnschild et al., 2010). This unique combination of powered flight, mammalian physiology, and complex vocal behavior has positioned bats as an emerging model system for studying a wide range of mammalian behaviors, including vocal learning, spatial navigation, and social behavior (Fernandez et al., 2021; Las & Ulanovsky, 2024; Palgi et al., 2025; Yartsev & Ulanovsky, 2013). Understanding how these complex behaviors emerged and acquired their present form requires considering the physiological and ecological demands that shaped them.

The integration of sound production with flight is highly conserved in bats, with decades of research showing that the emission of echolocation calls is tightly coupled to the wingbeat and respiratory cycles (Falk et al., 2015; Holderied & Von Helversen, 2003; Kalko, 1994; Koblitz et al., 2010; Schnitzler, 1971; Stidsholt et al., 2021; Suthers et al., 1972; Xia et al., 2025). This coupling is only briefly interrupted during prey capture (Stidsholt et al., 2021). Synchronization is generally considered energetically beneficial because the contraction of powerful muscles during flight supports the generation of high subglottal pressures needed for vocalizing (Lancaster et al., 1995; Speakman & Racey, 1991). Based on this mechanical linkage, it has been proposed that flight might have offered bats the unique opportunity to evolve more complex social vocalizations in the first place, thus explaining why elaborate songs are otherwise rare among mammals (Smotherman et al., 2016). First evidence supporting this comes from Spix’s disc-winged bats (*Thyroptera tricolor*), where the stationary emission of social calls leads to a 2-fold increase in metabolic rates (Chaverri et al., 2021).

However, despite the fact that many bat species emit some of their most complex vocalizations during flight (Smotherman et al., 2016), social calls are still frequently studied in isolation from flight (e.g. Babl et al., 2025; Burchardt et al., 2019; Chaverri et al., 2021). This is primarily due to the challenges of recreating the necessary social and environmental contexts in a laboratory setting, and the difficulty to capture high-quality synchronized recordings of flight and vocal behavior in the field (Jahelková et al., 2008).

Comparative research in birds, the only other flying vertebrate clade emitting highly complex song, has demonstrated that the evolutionary emergence of complex vocalizations is often associated with a relaxation of locomotor coupling: while non-vocal learners (species with largely innate song repertoires) typically maintain a 1:1 ratio between wingbeat and call emission, vocal learners have decoupled these systems, allowing for a more independent control of song (Berg et al., 2019). This raises a compelling question for bats: Complex song-like social communication is documented for species from five families (Smotherman et al., 2016), yet all bats retain a strong legacy of vocal-motor coupling for echolocation. It remains unknown whether the emission of complex social vocalizations during flight is similarly coupled by the underlying patterns of wingbeat and respiration, or if bats have decoupled these systems similar to birds to support the high degree of flexibility required for complex social communication.

To address this question, we investigated complex social vocalizations in wild Nathusius’ pipistrelles (*Pipistrellus nathusii*), a migratory bat whose complex social communication behavior is particularly well suited for examining vocal-motor coupling during flight. Males of this species produce a long, complex social call used for mate attraction and territorial defense, consisting of a stereotyped sequence of motifs that spans several hundred milliseconds, far exceeding a single wingbeat cycle of 90–100 ms (Holderied & Von Helversen, 2003; Jahelková et al., 2008; Kalko, 1994). As a result, each social call necessarily spans multiple wingbeats, much like aerial songs in vocal-learning birds (Berg et al., 2019; Smotherman et al., 2016).

*P. nathusii* also exhibits a tight coupling between the wingbeat and echolocation call (EC) rhythm, with a near 1:1 relationship (Holderied & Von Helversen, 2003; Kalko, 1994). This coupling is characteristic for all pipistrelles (Kalko & Schnitzler, 1993) and renders the EC pulse interval a reliable acoustic proxy for the underlying wingbeat period. Because social calls are embedded within sequences of regular echolocation calls during straight, steady flight, they can be unambiguously assigned to a single flying individual (Götze et al., 2020). Together, these species-specific properties create an ideal system in which the temporal structure of a complex social vocalization can be interpreted relative to the locomotor rhythm that constrains it.

Previous work suggests two mechanisms by which social calls might be integrated into the wingbeat cycle: (i) pre-planning based on the syllable content of individual motifs, as seen in echolocation call groups where call onsets shift to maintain energetically favorable timing in the wingbeat cycle (Falk et al., 2015; Koblitz et al., 2010; Xia et al., 2025), and (ii) a rhythmic-scaling mechanism, in which motif onsets follow a rhythmic temporal structure proportional to the respiratory or wingbeat cycle, as indicated by the isochronous rhythm found in territorial social calls of other bats (Burchardt et al., 2019).

Here, we combined audio recordings and synchronized video recordings collected in the field over multiple years with a kinematic inference model to determine how *P. nathusii* integrates its complex social calls into flight. We found that motif durations scaled with syllable number, but motif onsets were governed by a rhythmic process anchored to the wingbeat cycle. The entire social call was precisely integrated into three consecutive wingbeats, with motifs emitted at narrow, preferential phases of the wingbeat cycle. This represents the first evidence for how complex social calls are integrated into the wingbeat mechanics of bat flight, revealing that vocal-motor coupling extends far beyond echolocation to shape the structure of social communication.

## Materials and Methods

### Data collection

We recorded wild, flying bats of the species *Pipistrellus nathusii* over the course of multiple years. All recordings were conducted at the Federseemuseum in Bad Buchau, Germany (48.069723° N, 9.611259° E).

Data collection was split into two phases: an initial audio-only phase, with recordings collected over five nights in late May, 2022. A subset of 2.5 h of audio recordings from this period was used in this study. The later synchronized video and audio recordings were collected over five days in May 2024 and 2025, totaling over 11 hours of recording time.

### Recording setup

For initial audio recordings, the setup consisted of a microphone mounted on a tripod at a height of 1.35 m above the ground. For later sessions with combined video recordings, a camera was mounted 18 cm next to the microphone. In 2022 and 2024, devices were set up in front of a known roost of *P. nathusii* located in the building’s outer wall at a horizontal distance of 8.6 m. In 2025, the setup was moved on one occasion to the other side of the building at a new roost location, at a horizontal distance of 6.4 m.

For audio recordings we used the PC-Tape recording system (Animal Physiology, University of Tübingen, Germany) with a custom-built ultrasonic microphone (almost flat frequency response at 100 kHz, 4 dB less sensitive than at 20 kHz) and a sampling frequency of 480 kHz (Melcón et al., 2007). A post trigger stored the content in a ring buffer, allowing it to manually save audio recordings of custom duration retrospectively. For the initial audio-only phase, 90 samples of 5 s each were recorded after social call emission was detected.

Video recordings utilized a digital video camera with an effective temporal resolution of 50 Hz (25 interlaced frames^-s^ or 50 half-frames^-s^). An infrared strobe system provided 1 ms flashes at 50 Hz, synchronized with the camera’s field rate, thus each half-frame was illuminated by a separate strobe flash (Koblitz et al., 2010; Sändig et al., 2014). Recordings were stored analogously (Sony DCR-PC8E camcorder) and digitalized using the Simi-Motion software (Simi Motion 6.5, SIMI Reality Motion Systems GmbH).

For all audio-video coupled recordings, continuous video was recorded each night whenever bat activity was present, accumulating over 7 hours of video footage analyzed. Precise synchronization of the microphone and camera was achieved using the VITC-code of the PC-tape, resulting in a synchronization accuracy of ± 1 ms. Synchronized audio recordings were saved retrospectively whenever a bat passed through the camera’s field of view, yielding 910 audio samples of various length, totaling 5 hours and 3 minutes of audio data analyzed.

All audio samples were manually screened for social calls. Video recordings were checked for clear, high-quality recordings of bats during straight search flight (i.e. absence of flight maneuvers like sharp turns or loops) or while emitting social calls. All recordings were conducted in accordance with relevant guidelines and regulations.

### Social call analysis

Sound recordings were visualized as spectrograms using custom-made software Selena (University of Tübingen, Germany), with a temporal resolution of 0.06 ms and a spectral resolution of 156.3 Hz (FFT 512, Blackman window, 70 dB dynamic range).

Echolocation calls (ECs) were identified by their species-specific terminal frequency centered around 39 kHz (range 36–44 kHz; Zsebők & Görföl, 2012). Analyzed social calls consisted of ABC motifs, unique to our focal species (Fig. 1a, b; Furmankiewicz, 2003; Jahelková, 2011; Jahelková et al., 2008; Russ et al., 1998; Russ & Racey, 2007).

**Figure 1.**
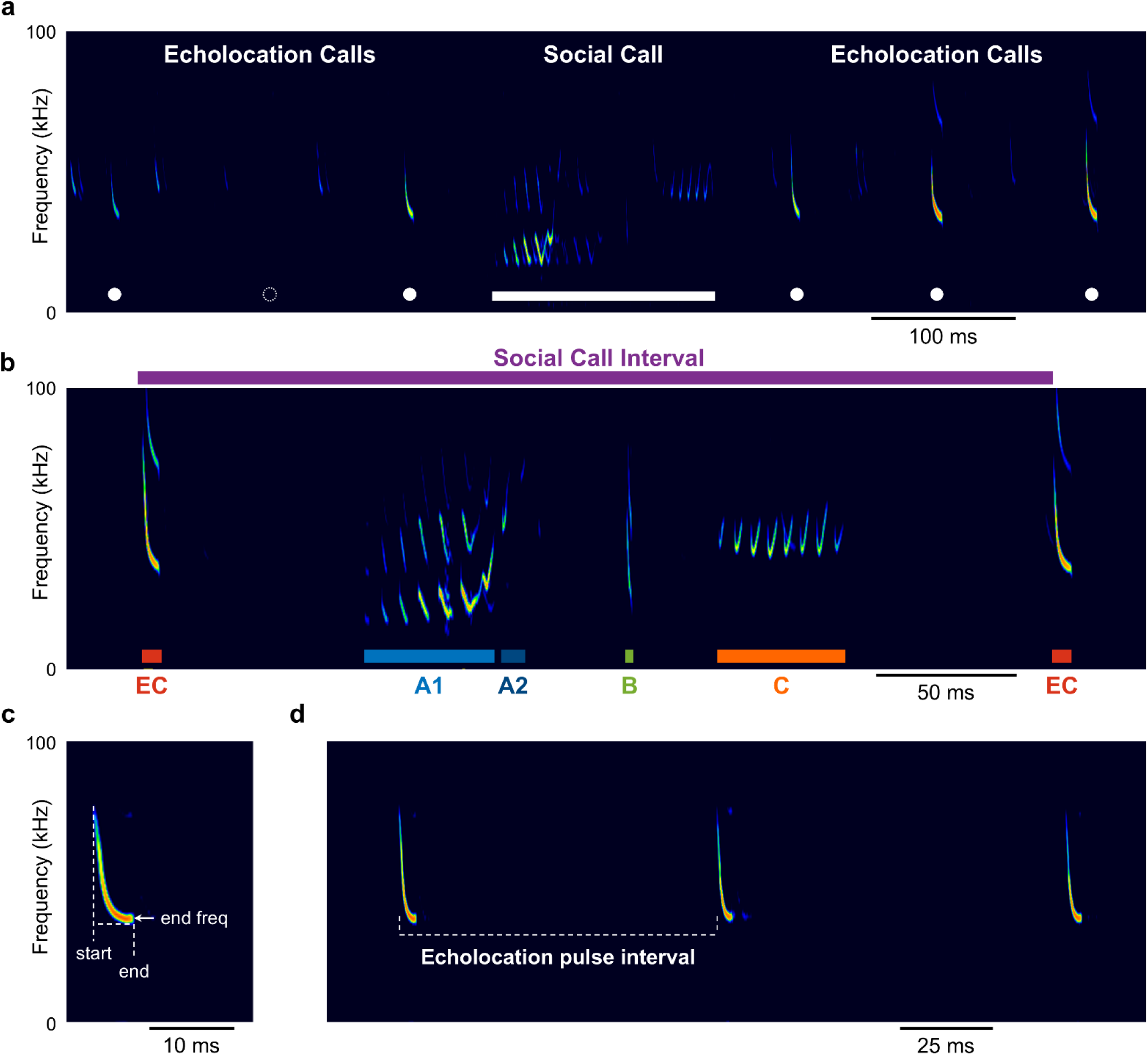
Social calls are embedded into a regular sequence of echolocation calls during flight. **a** Social call (solid line) flanked by echolocation calls (filled circles) of the same individual during flight. Before the social call, a single echolocation call is omitted (dotted circle). **b** Social call interval (SI) includes the time from the beginning of the echolocation call preceding the social call to the beginning of the echolocation call following the SC. A typical social call consists of the A1, A2, B and C motif. **c, d** Measured echolocation call parameters included start and end of each call, terminal frequency (**c**), and the pulse interval (PI) between consecutive calls (**d**).

To ensure social calls in the audio-only data were emitted during flight, we utilized the stability of acoustic features in the echolocation call sequence, which serves as a reliable indicator of individual flight (Götze et al., 2020). Therefore, we accepted only those social calls that were unambiguously embedded in a sequence of echolocation calls, resuming at an almost identical terminal frequency after the social call had ceased (Fig. S1a; before social call: 39.17 ± 0.51 kHz, after social call: 39.17 ± 0.46 kHz, p = 0.96, n = 37, paired t-test). Acceptance further required all main social call motifs (A1, B, C) to be clearly identifiable and measurable, with 25 out of 37 recordings also exhibiting the A2 sub-motif (67.57%; Jahelková et al., 2008). A total of 37 social calls were analyzed. The “social call interval” (SI) (Fig. 1b) was defined as the time period between the onset of the last EC before the social call and the onset of the first EC immediately following the social call.

Motif parameters (onset, duration, inter-motif interval, syllable number, start and terminal frequency) were measured manually based on spectrograms. Inter-syllable intervals were calculated for all multi-syllable motifs. We also measured the start and terminal frequency, duration and pulse interval of up to three ECs flanking each social call (Fig. 1c, d), totaling 136 analyzed ECs. The beginning and end of signals were defined as 14 dB relative to ambient noise levels.

The echolocation pulse interval (PI) was utilized as an acoustic proxy for the underlying wingbeat frequency during flight due to the species’ known 1:1 coupling of wingbeats and echolocation calls (Holderied & Von Helversen, 2003; Kalko, 1994; Schnitzler, 1971). Since *P. nathusii* occasionally emits only one EC per two wingbeats (Holderied & Von Helversen, 2003; Kalko, 1994), we observed a bimodal distribution of PIs (Fig. S2). To correct for this, a Gaussian Mixture Model was fitted to the distribution of all PIs. The model was constrained to yield two best-fitting Gaussians, ensuring neither distribution contained fewer than 10 samples. Out of 1000 runs, the most frequent crossover point distinguishing between single wingbeat and double wingbeat PIs was found at 130.07 ms (maximum range: 120.11–130.39 ms). This value served as the threshold to discriminate between call emissions at single and double wingbeats, with all PIs belonging to the double wingbeat distribution divided by two to make them comparable to single wingbeat call intervals. This threshold aligns well with previously reported values for the species (∼140 ms; Holderied & Von Helversen, 2003; Kalko, 1994) and was later confirmed using synchronized video recordings (max. PI for 1 call/wingbeat: 123.0 ms).

### Wingbeat reconstruction and mapping of calls

Recorded videos were analyzed using the Simi-Motion software (Simi Motion 6.5, SIMI Reality Motion Systems GmbH). Wingbeat reconstruction was performed on 40 recordings of straight flights in which the wing position could be accurately determined. To analyze the wing movements, the position of the wing’s tip was measured relative to the animal’s body midpoint in each frame and normalized (to a range of ±1). The 50 Hz temporal resolution of the video recordings was suitable for the expected wingbeat frequencies of 8–15 Hz in *P. nathusii* (Holderied & Von Helversen, 2003; Kalko, 1994)

The wingbeat cycle was reconstructed by fitting a sinusoidal function to the measured relative wing tip positions. This was achieved using MATLAB’s fit function with a custom fit-type, described by the equation:

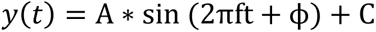

where y(t) is the normalized wing tip position at time t, and fitting parameters were amplitude (A), frequency (f, wingbeat frequency), phase shift (ϕ), and vertical offset (C). This approach yielded a high average goodness-of-fit for echolocation call sequences (R^2^ = 0.89 ± 0.01). To also incorporate echolocation calls emitted closely before or after the tracked video sequences, reconstructed flightpaths were extrapolated by ±50 ms. Additionally, call onset times were corrected for audio delay over distance based on the animal distance and the speed of sound at 20°C air temperature (343.2 m/s). For this, recording sites were mapped each night based on field of view of the camera and horizontal and vertical distances to the animal were subsequently estimated by two independent, blind annotators.

Echolocation call onsets were mapped directly onto the reconstructed sinusoidal wingbeat cycle. The angle of the sinusoidal function at each time point was used as the phase of the wingbeat (0° to 360°). In this framework, 0° to 180° represents the downstroke, and 180° to 360° represents the upstroke. Following this approach, a total of 88 calls were mapped onto the underlying wingbeat cycle (see Fig. 4a, b).

For empirical validation of our model’s predictions, three individuals (indicated by differences in EC terminal frequencies) were tracked while actively emitting a social call during straight flight. These tracked video recordings covered a total duration of 200–380 ms, resulting in 2 to 4 fully reconstructed wingbeats. However, reconstructed flight segments only partially covered the full duration of the social call emission. As only straight flight trajectories were included in the analyses, flight paths were extrapolated accordingly to cover the full SI duration, with tracked portions of the SI duration varying substantially (69%, 36%, and 16%). Reconstructed sinusoidals exhibited a good overall fit (R^2^ = 0.61 ± 0.13).

### Monte Carlo inference model

A Monte-Carlo simulation consisting of 10,000 runs was used to estimate the timing of social call motifs relative to the wingbeat cycle from audio-only recordings. This simulation was essential because the acoustic data provide the relative temporal structure of the social call, but do not contain information about its absolute position within the wingbeat cycle.

The simulation framework was based on three empirically derived observations. (1) The temporal intervals between motifs (EC→A1, A1→A2, A2→B, B→C, C→EC) can be expressed in units of wingbeats using each individual’s pulse interval (PI), yielding their relative spacing even in the absence of wingbeat measurements. (2) Social calls are seamlessly embedded within the ongoing echolocation call (EC) sequence, and the duration of the SI reliably corresponds to approximately three wingbeats, indicating that SI onset and offset are anchored to the underlying wingbeat rhythm. (3) Video tracking demonstrated that the EC preceding the SI is emitted at a characteristic phase of the wingbeat cycle. Thus, the dominant source of uncertainty in audio-only sequences is the exact wingbeat phase of this initial EC. The simulation could therefore propagate this uncertainty while preserving the temporal structure among motifs.

For each iteration, the starting EC phase was randomly drawn from a normal distribution derived from our video-tracked dataset (mean = 315.87°, SD = 47.08°, N = 40). A set of motif intervals and durations from a single audio recording was then randomly selected from the audio-only dataset (with replacement), and used for that run. To preserve the natural variability in motif composition, each run also included a random decision about whether to simulate an A2 motif, based on its empirical occurrence probability (25/37 files; 67.6%). Although the interval between A1 and B does not differ significantly between A2-present and A2-absent cases (A1→B = 1.11 wingbeats vs. A1→A2→B = 1.08 wingbeats; p = 0.6055, two-sample t-test), incorporating this branching structure allowed the simulation to reflect the observed variation in motif sequences.

Motif intervals and durations were expressed as multiples of the individual wingbeat period (as approximated by each individual’s pulse interval), which specifies their relative timing with respect to the starting EC. These relative timings were then aligned to the randomly sampled starting EC phase to determine motif onsets in degrees, thereby preserving the empirically measured motif structure while systematically propagating uncertainty in their absolute placement within the wingbeat cycle. Finally, motif offsets were calculated by adding motif durations to their respective onset times.

Simulation outputs (motif onsets and motif centers) were subsequently tested for significant circular clustering using a bootstrap-based circular statistics approach described below. Based on the simulated motif onsets and offsets, we also computed the normalized probability of each motif being present at any given wingbeat angle in steps of 1°, and binned these probabilities into three equal intervals. For visualization, motif onsets, durations, and syllable counts were mapped onto a sinusoidal representation of the wingbeat cycle using the simulation means, accompanied by standard deviations from the simulation and error-propagated SEMs derived from the real data via RSS (n = 37).

### Statistics

All statistical analyses were conducted using custom-written MATLAB (Version R2024b) and Python (Version 3.11.10) software. Prior to each analysis, normality was assessed using the Shapiro–Wilk test.

Linear Mixed-Effects Models were utilized to analyze motif onset timing. These models included the respective predictor variable, motif type, and their interaction as fixed effects, and a random intercept per file to account for repeated measures (i.e., each recording contributes multiple motif onsets – A1, A2, B, and C – that share the same predictor value) and unbalanced motif occurrence (A2 presence/absence). Pairwise mixed-effects models were used to compare group means between motifs while accounting for repeated observations.

All other linear relationships were examined by fitting separate linear regression models to estimate the regression slope, 95% confidence intervals, p-value, and R^2^.

Paired t-tests were employed to compare EC variables within each recording, including the interval and terminal frequency before and after the SI. Pairwise t-tests (Welch) with Bonferroni correction were used to compare inter-syllable intervals of multi-syllable motifs. The SI/EC relationship was tested using a one-sample t-test against the theoretical integer value of 3.

The synchronization of the wingbeat and echolocation calls was assessed using the MATLAB CircStat Toolbox (Berens, 2009). Using its circular statistics functions, mean call position and variability were calculated for each individual recording. For the final population analysis, we used the mean of each recording as an independent sample. We used the Rayleigh test included in the toolbox to check if the circular distribution of mean EC onsets (0 to 360°) was significantly different from a uniform distribution.

To evaluate the circular predictions of the Monte-Carlo inference model (motif onsets and centers), the standard Rayleigh Test was deemed unsuitable, as it is highly sensitive to the extremely large sample size (10,000 iterations). Therefore, a custom bootstrap test and respective empirical p-values were calculated to provide results independent of the number of Monte Carlo simulation runs. This test determined if the circular clustering observed in our simulated data was significantly different from a uniform distribution (0° to 360°) across 10,000 bootstrap iterations. In each iteration, 200 samples of simulated motif onsets and centers, respectively, were subsampled without replacement and compared against the same number of samples drawn from a uniform null distribution. The circular vector strength (R), used to accurately assess the mean of circular data, was calculated for both the simulated and null subsampled data using the CircStat Toolbox in each iteration. Finally, the empirical p-value (one-sided) was defined as the frequency with which the circular vector length of the null data (R_null_) was larger than or equal to the circular vector length of the simulated data (R_siml_).

All reported values represent the average ± standard error of the mean if not stated otherwise. The sample size of video tracked social calls (n=3) was too small for robust statistical analysis; observed motif phases were therefore compared qualitatively against the inference model’s predictions.

## Results

### Complex social calls follow a rhythmic temporal structure scaled to the overall call interval

We investigated the temporal structure and integration of complex social calls (SCs) during free flight in wild *P. nathusii*. SCs were analyzed from audio recordings only when they occurred midflight, defined by the interruption and subsequent resumption of a regular echolocation call (EC) sequence, indicative of steady flight in pipistrelles (Fig. 1a, b; Götze et al., 2020). The acoustic parameters of the flanking ECs (Fig. 1c, d) remained highly consistent, with the average terminal frequency and pulse interval remaining similar before and after the SC (Fig. S1a, b; Terminal frequency before: 39.17 ± 0.51 kHz, after: 39.17 ± 0.46 kHz, p = 0.96, n = 37, paired t-test; Pulse interval before: 94.52 ± 2.27 ms, after: 91.95 ± 2.47 ms, p = 0.44, n = 25, paired t-test). This indicated that the same bat that had produced the initial echolocation sequence resumed emitting regular orientation calls after its midflight social call. Thus, the echolocation calls flanking the SC provided us with a stable reference point for all SCs emitted during flight. Subsequently, the period between the onset of the preceding EC and the onset of the succeeding EC was defined as the Social Call Interval (SI; Fig. 1b), with an average duration of 283.65 ± 5.03 ms. Each SI contains four distinct motifs, three multi-syllable motifs (A1, A2, C) and one single-syllable motif (B).

We first hypothesized that motif timing was pre-planned based on each motif’s syllable content, similar to timing adjustments observed in bat echolocation call groups, where call onsets shift based on the number of calls produced per group (Falk et al., 2015; Koblitz et al., 2010; Xia et al., 2025). These shifts are thought to keep the call group centered around the energetically most viable phase of the wingbeat cycle. If social calls were structured in the same way, this hypothesis predicts that motifs with more syllables should be initiated earlier relative to the start of the SI. We examined whether motif onset times correlated with the variable number of syllables produced in the respective multi-syllable motifs. Surprisingly, onset times for all three multi-syllable motifs were not systematically correlated with the number of syllables produced (Fig. 2a; A1: R² = 0.003, p = 0.76; A2: R² = 0.011, p = 0.61; C: R² = 0.002, p = 0.79; LM). Furthermore, the onset of motifs B or C did not shift systematically with the syllable count of the preceding motif (Supp. Fig. ##; B: R^2^ = 0.065, p = 0.13; C: R^2^ = 0.083, p = 0.08). Instead, the overall duration of each motif scaled almost perfectly with its syllable count (Fig. 2b; A1: R² = 0.90, p < 10^-6^; A2: R² = 0.95, p < 10^-6^; C: R² = 0.94, p < 10^-6^), suggesting highly regular inter-syllable intervals (Fig. 2c).

**Figure 2.**
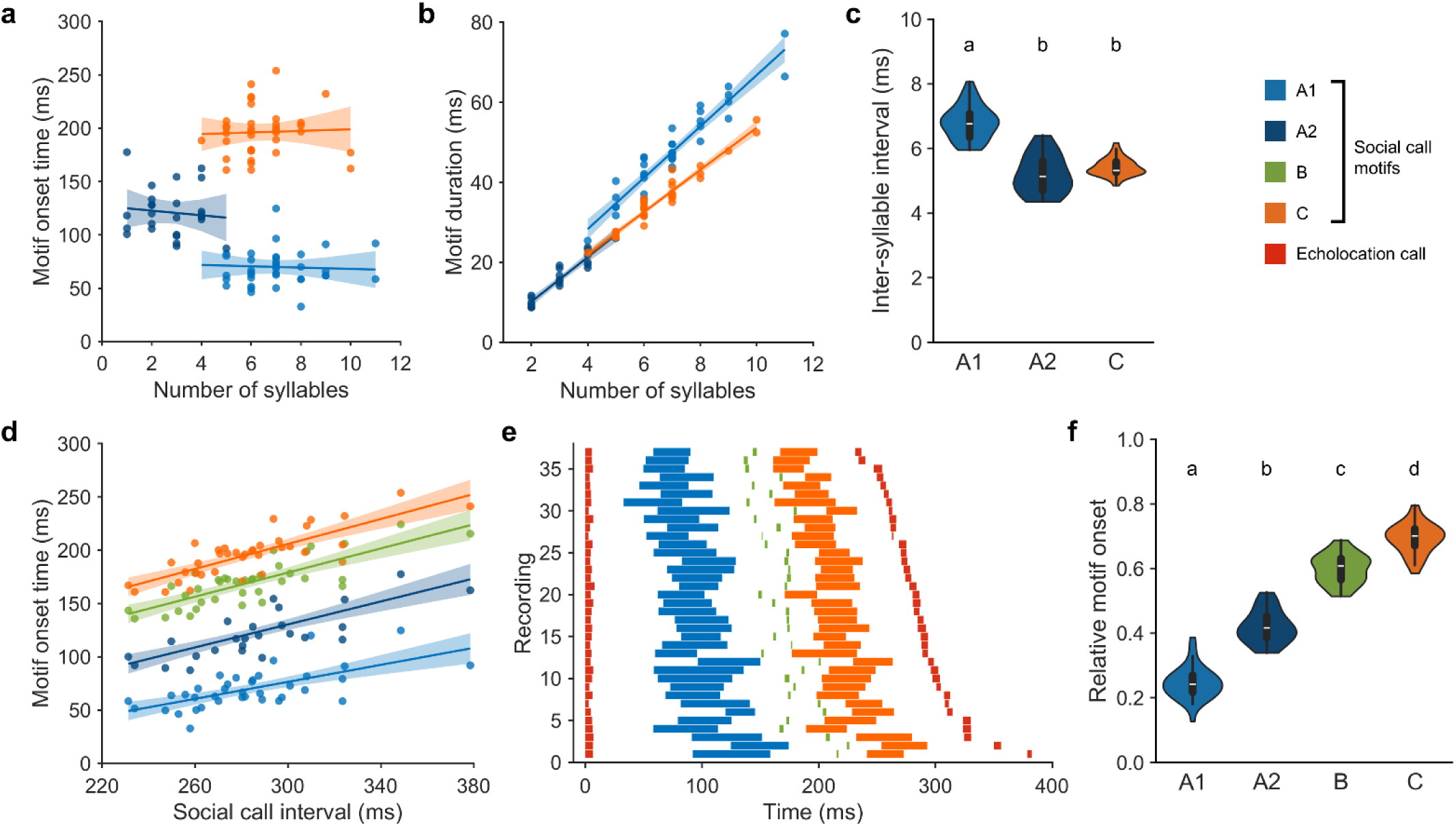
Motif onsets scale relative to the duration of the social call interval. **a, b, c** Motif onset time does not depend on number of syllables in each multi-syllable motif (**a**), instead, number of syllables determines motif duration (**b**), with each motif exhibiting stereotyped inter-syllable intervals (**c**). **d** Motif onset times depend on absolute duration of the social call interval, with onset times shifting to later time points for longer social call intervals. **e** Onsets and offsets of all elements of the social call interval plotted over time for each recording, sorted by absolute SI duration. **f** Shift in onset times results in stereotyped motif onsets relative to respective social call interval. Distinct letters above violins denote significant differences between groups after Bonferroni correction.

The absence of adaptive pre-planning based on motif content suggests that motif onset timing is governed by a temporal process independent of each motif’s internal acoustic structure. This raised the possibility that the sequence of motifs follows a simple rhythmic framework, a kind of internal clock, that scales motif onsets relative to the overall duration of the social call interval.

To test this relative-scaling hypothesis, we assessed if the onsets of all motifs scaled proportionally with the overall SI duration. We found strong positive correlations between the onset time of each motif and the respective SI duration across individuals (Fig. 2d, e; A1: R² = 0.43, p < 10^-6^; A2: R² = 0.62, p < 0.005; B: R² = 0.64, p < 0.001; C: R² = 0.65, p < 10^-3^). This proportionality indicates that the emission of each motif maintains a relatively stable temporal position within the SI, regardless of the SI’s absolute length. When expressed relative to the overall SI duration, motif onsets clustered into narrow, distinct intervals: the A1 motif began at 25%, A2 at 42%, B at 60%, and C at 70% of the total SI duration (Fig. 2f, p < 10^-4^ for all pairwise comparisons, Bonferroni corrected).

Collectively, these results demonstrate that while motif duration is dependent on syllable count, motif onset is rigidly controlled by a rhythmic process that scales with the overall SI duration. This raises the question which external factor scales the overall SI duration, while maintaining stable proportional timing of the motifs within that interval.

### The social call interval is integrated into the wingbeat cycle

Given that motif timing followed a rhythm scaled to the overall SI duration, we investigated which factor governed SI length. It is well known that bats tightly synchronize echolocation calls (ECs) with their respiratory and wingbeat cycles during flight, often adhering to a 1:1 coupling ratio (Kalko, 1994; Schnitzler, 1971; Suthers et al., 1972). We therefore hypothesized that the SI duration is mechanically constrained by the wingbeat rhythm. To test this, we first measured the pulse interval (PI) in regular echolocation sequences (Fig. 1d) revealing a mean pulse interval of 91.47 ± 1.21 ms. Since Nathusius’ pipistrelles typically emit one EC per wingbeat, the PI serves as a reliable approximation of the individual wingbeat period during straight flight. This ∼90 ms period aligns well with prior reports for this species (Holderied & Von Helversen, 2003; Kalko, 1994) and validated our approach of approximating flight kinematics purely from audio recordings.

To determine if the SI duration is governed by the underlying wingbeat rhythm, we first correlated SI duration with each individual’s wingbeat period (as approximated by the PI). We found a significant positive relationship between SI duration and PI across individuals (R = 0.39, p < 0.005; Fig. 3a), demonstrating that longer wingbeat periods resulted in proportionally longer SI durations. This result suggests that the SI temporal boundaries are subject to motor constraints imposed by the flight cycle.

**Figure 3.**
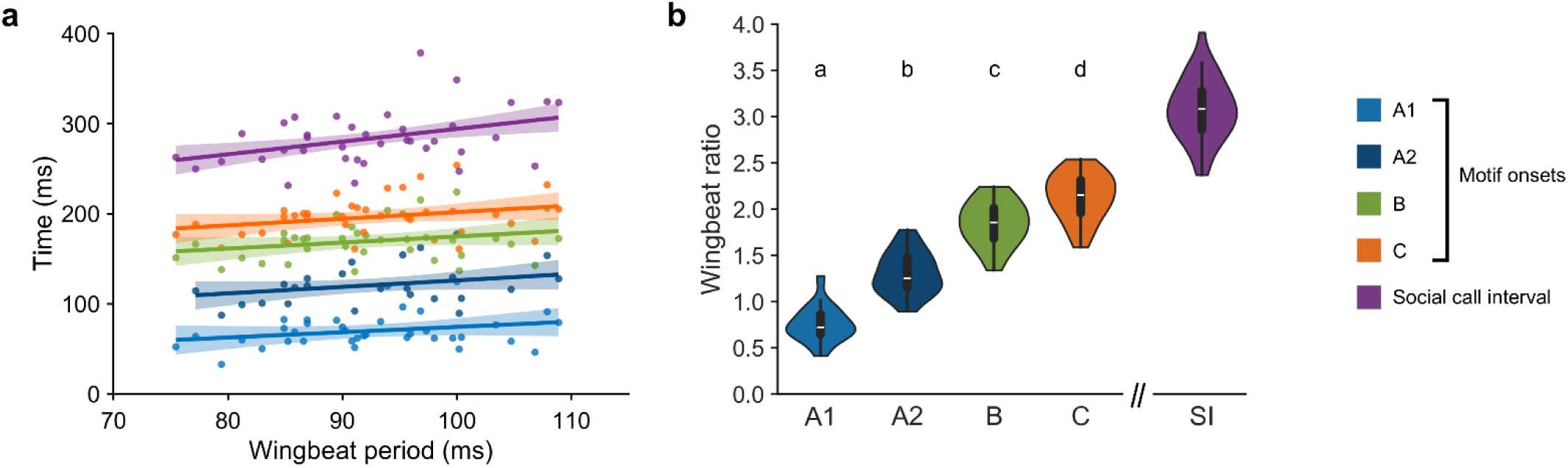
Social call interval is integrated into the wingbeat cycle. **a** Social call interval duration correlates significantly with wingbeat period of individuals. Motif onsets show a weak correlation with wingbeat period. **b** Motif onsets occur after a distinct, stereotyped number of wingbeats within the social call (left). The overall social call interval encompasses three full wingbeats on average (right). Distinct letters above violins denote significant differences between groups after Bonferroni correction.

Based on the observed synchronization, we hypothesized that the SI duration should represent an integer multiple of the wingbeat period. This prediction stems from the necessity to integrate the social call within the continuous 1:1 wingbeat to call cycle, allowing the rhythmic EC sequence to resume seamlessly upon the SI’s completion. Consistent with this idea, the pulse interval of ECs immediately before and after the SI did not differ significantly (Fig. S1c; before social call: 93.01 ± 2.87 ms, after social call: 91.88 ± 3.94 ms, p = 0.79, n = 24, paired t-test), indicating that the call sequence resumed without any temporal adjustment.

By calculating the ratio of each bat’s individual SI duration relative to its wingbeat period, we found that the mean ratio clustered tightly around 3:1 (mean ratio: 3.08 ± 0.05; t-test against 3: p = 0.1550, n=37; Fig. 3b). This finding strongly suggests that the social call interval consistently and precisely accommodates three complete wingbeats between the preceding and succeeding echolocation calls, revealing a highly rigid vocal-motor coupling.

Finally, we tested how the motif onsets are expressed relative to the wingbeat cycle. Motif onsets showed only weak to moderate correlations with the absolute wingbeat period (A1: R = 0.27; A2: R = 0.39; B: R = 0.26; C: R = 0.28; p > 0.05 for all; Fig. 3a). However, when expressed relative to the wingbeat period, the onsets clustered narrowly around fixed wingbeat ratios across individuals (Fig. 3b). Specifically, motif A1 occurred on average after 0.76 ± 0.03 wingbeats, A2 at 1.31 ± 0.05 wingbeats, B at 1.84 ± 0.04 wingbeats, and C at 2.13 ± 0.04 wingbeats (p < 10^-5^ for all pairwise comparisons, Bonferroni corrected).

These findings demonstrate that the wingbeat cycle governs the duration of the entire social call interval, revealing a precise 3:1 vocal-locomotor coupling. Within this coupling, the onset of individual social call motifs is dynamically scaled to maintain emission at narrowly defined, preferential phases of the underlying wingbeat cycle.

### Echolocation call emission in the wild is coupled to the upstroke phase of the wingbeat

To lay the empirical groundwork for our subsequent mechanistic inference model of vocal–motor coupling, we first aimed to quantitatively validate the assumptions underlying the audio-based analysis. To this end, we performed synchronized video and audio recordings of wild *P. nathusii* during free flight. To our knowledge, this represents the first quantitative characterization of wingbeat–echolocation synchronization in wild individuals of this species. This approach allowed us to reconstruct the wingbeat cycle and align it with emitted echolocation calls.

The mean wingbeat period measured was 88.48 ± 1.05 ms, which closely matched the approximated wingbeat period derived solely from the EC pulse intervals of 91.47 ± 1.21 ms (p = 0.40, two-sample t-test, n = 40). This confirmed the reliability of the PI as a proxy for the wingbeat period in this species. We found that usually a single EC was produced during each wingbeat, confirming a 1:1 coupling ratio (Fig. 4a, b). Pipistrelles sometimes exhibit wingbeats without call emission. The frequency of single ECs being emitted across two wingbeats of 9.1% (8/88 of all tracked ECs), matched both previous reports (Holderied & Von Helversen, 2003) and our earlier analysis of the audio-only recordings (13.5%; 18/133 calls). Combined, these video-tracking results verified the fundamental synchronization between wingbeat and echolocation calls in *P. nathusii*, therefore confirming earlier auditory analyses and providing a prerequisite for the subsequent mechanistic modeling.

**Figure 4.**
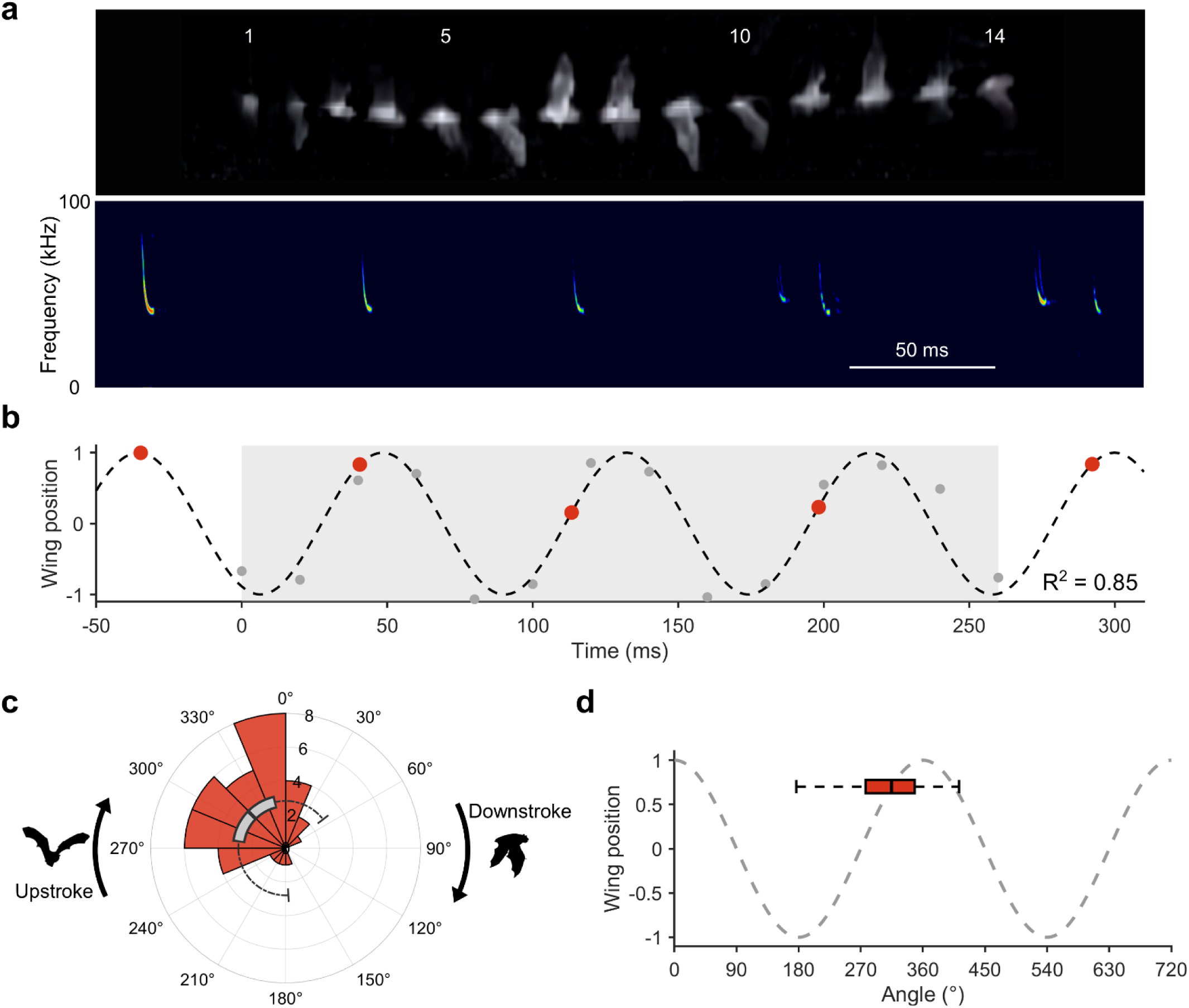
Echolocation call emission in the wild is coupled to the upstroke phase of the wingbeat. **a** Illustration of exemplary data collected from an individual flying bat in the field. Upper illustration shows stacked images of a video recording used to reconstruct the wingbeat cycle, with corresponding frame numbers shown above selected frames. Frames are roughly aligned to the spectrogram below. At the end of the recording, a second bat of a different species enters the recording area as indicated by two calls appearing with at a new terminal frequency. **b** Reconstructed wingbeat cycle of the animal shown in **a**. Sinusoidal fitted to wing position measurements (grey circles) with R^2^ = 0.85. Echolocation calls (red circles) plotted at time delay-corrected positions on the wingbeat cycle. Grey shaded area denotes the time interval covered by the video recording. **c** Polar histogram of population data, showing each individual’s mean wing angle at which echolocation calls were emitted (n = 40). 0°–180° denotes downstroke, 180°–360° denotes upstroke phase of the wingbeat. Circular boxplot of population data, showing median, IQR (grey box) and full range (whiskers) of wing angles at which echolocation calls were emitted. Note, no call emission during the latter half of the downstroke phase. **d** Same boxplot of population call emission data as in **c** aligned to the wingbeat cycle.

Crucially, we quantified the specific phase of the wingbeat cycle during which regular ECs were emitted across individuals. We found that ECs were almost exclusively produced during a tight window at the end of the upstroke phase, preceding the transition to the downstroke (315.87 ± 7.44°, n = 40; Fig. 4c, d). This emission timing was highly stereotyped (p < 10^-8^, Rayleigh Test; 95% CI: 298.66–333.08°), demonstrating precise vocal-motor coupling. Notably, a large segment of the downstroke phase (52°–177°) was essentially devoid of EC emission (Fig. 4c). Results did not differ when using all individually tracked calls (Fig. S3; Mean call phase = 320.71 ± 4.52°, n = 128).

Our video tracking validated the fundamental assumption of our audio-based analysis that the EC-to-wingbeat coupling is 1:1. Most importantly, it confirmed that the PI serves as a reliable proxy for the wingbeat period, and that the emission of ECs occurs at a highly stereotyped phase within the wingbeat cycle. Combined with the 3:1 SI-to-wingbeat ratio and the rhythmic process governing motif onsets, these parameters enabled the construction of a simple mechanistic model to predict how the social call’s temporal structure aligns with the underlying wingbeat cycle.

### Inference model predicts motif emission at specific wingbeat phases

To mechanistically map the entire social call structure onto the underlying wingbeat cycle, we integrated three core constraints validated in earlier sections into a Monte Carlo inference model. These constraints were: 1) motif onsets maintain stable relative timing within the SI, while durations are governed by syllable count; 2) social calls are seamlessly embedded within the ongoing EC sequence, scaled to span three wingbeats; and 3) the flanking ECs are emitted at a highly stereotyped wingbeat phase. Using this model, we predicted the precise phase of the wingbeat at which the onsets and offsets of all four social call motifs should occur during flight (see Methods for more details).

The inference model revealed that each motif is predicted to occur within a distinct and specific phase of the three-wingbeat cycle (Fig. 5a). Specifically, the A1 motif is predicted to start near the turning point into the first upstroke (229.13°), primarily spanning the latter two-thirds of the upstroke phase. The subsequent A2 motif follows immediately (onset at 63.41°), spanning across most of the first downstroke phase. Emission of the single-syllable B motif is inferred to be around the center of the second upstroke (255.86°), and the C motif onset was predicted to coincide with the turning point into the third downstroke (359.63°), covering the majority of this final downstroke phase. All motif onsets were significantly coupled to specific wingbeat phases (Fig. 5b; A1: p < 10⁻³, A2: p = 0.0015, B: p = 0.0133, C: p = 0.0149; bootstrap test, n = 200).

**Figure 5.**
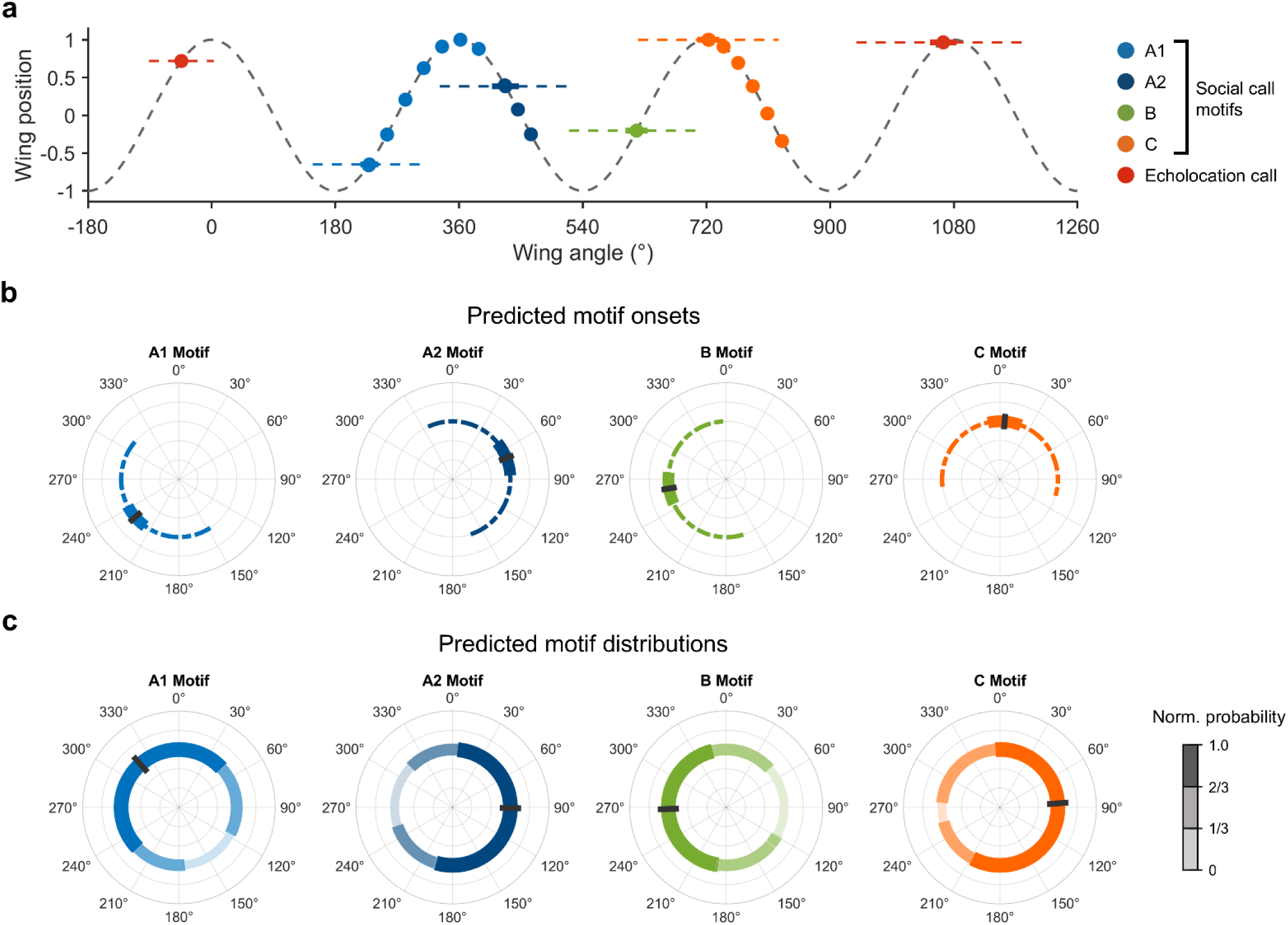
Inference model predicts motif emission at specific wingbeat phases. **a** Mean onsets, durations and syllable numbers of all elements of the social call interval mapped onto the wingbeat cycle as predicted by the Monte Carlo simulation. Each element onset is shown with error-propagated SEM derived from real data (thick horizontal bars) and standard deviation of the simulation (dashed horizontal bars). **b** Circular plots showing predicted motif onsets relative to wingbeat phase. For each motif, mean wingbeat phase at onsets (black vertical bars), SEM (thick horizontal bars) and standard deviation of onsets (dashed horizontal bars) are shown. **c** Circular plots showing predicted distribution of full motifs relative to wingbeat phase. For each motif, normalized probability of the respective motif being present at any given wingbeat phase are shown with mean center of the distribution (black vertical bars). 0°–180° denotes downstroke, 180°–360° denotes upstroke phase of the wingbeat.

Although motif durations vary widely with syllable number (s. Fig. 2b), the predicted distribution of full motif durations was still restricted to certain wingbeat phases (Fig. 5c), with motif centers significantly coupled to distinct wingbeat phases (A1: 321.91°, p < 10⁻³, A2: 90.81°, p = 0.0053, B: 260.74°, p = 0.0122, C: 63.31°, p = 0.012; bootstrap test, n = 200). Consequently, all motifs occupy characteristic, though less sharply defined wingbeat phases. This pattern suggests that social call emission remains phase-locked to the wingbeat, even with flexible motif durations.

The model further predicted the timing of the closing EC of the SI, which marks the transition back to regular echolocation sequence. This EC is predicted to occur at the end of the third wingbeat at approximately 335.98° during the upstroke phase. This angle is highly similar to the starting phase determined for regular EC sequences (315.87°), suggesting that the three-wingbeat interval precisely accommodates the social call while ensuring the resumption of the regular EC sequence remains energetically and kinematically advantageous.

The inference model provides a mechanistic explanation for the observed scaling patterns in the acoustic data, generating a testable hypothesis: each motif of the social call is preferentially emitted at a specific phase of the underlying wingbeat cycle, constrained within a fixed three-wingbeat interval. The SI stretches or shortens based on the underlying wingbeat period, in order to smoothly integrate the social call into the EC sequence during flight and enable motif emission at stereotyped wingbeat phases.

### Video-tracked social calls confirm model predictions of stereotyped motif emission

To empirically test the predictions of our inference model, we analyzed synchronized video and audio recordings of three individuals emitting complex social calls during straight flight (mean wingbeat period: 83.33 ± 3.34 ms). As direct tracking could only cover part of each social call interval, the remaining phases were extrapolated in straight-flight segments where wingbeat dynamics are highly stereotyped. Using this approach, we mapped the social call and adjacent echolocation calls onto the wingbeat cycle of each individual.

Overall, the observed temporal structure of the tracked social calls closely resembled the predicted patterns (Fig. 6), confirming several key predictions of the inference model:

1. All ECs flanking the SI were emitted during the upstroke phase (range: 245.10°–308.71°), consistent with our findings from regular EC sequences. 2) The A1 motif onset occurred around the turning point from downstroke to upstroke of the second wingbeat in two out of three individuals (144.14° and 154.20°). 3) The single A2 motif observed in these recordings appeared immediately after A1 had ceased, right after the turn point into the second downstroke (2.71° wingbeat angle), landing well within the predicted interval. 4) The B motif consistently occurred during the first half of the upstroke (range: 204.60°–228.86°), aligning perfectly with our predicted interval. 5) The onset of the C motif in all three individuals occurred at the predicted phase, specifically right before the transition into the third downstroke phase (range: 311.08°–343.91°), closely matching the model’s mechanistic prediction.

**Figure 6.**
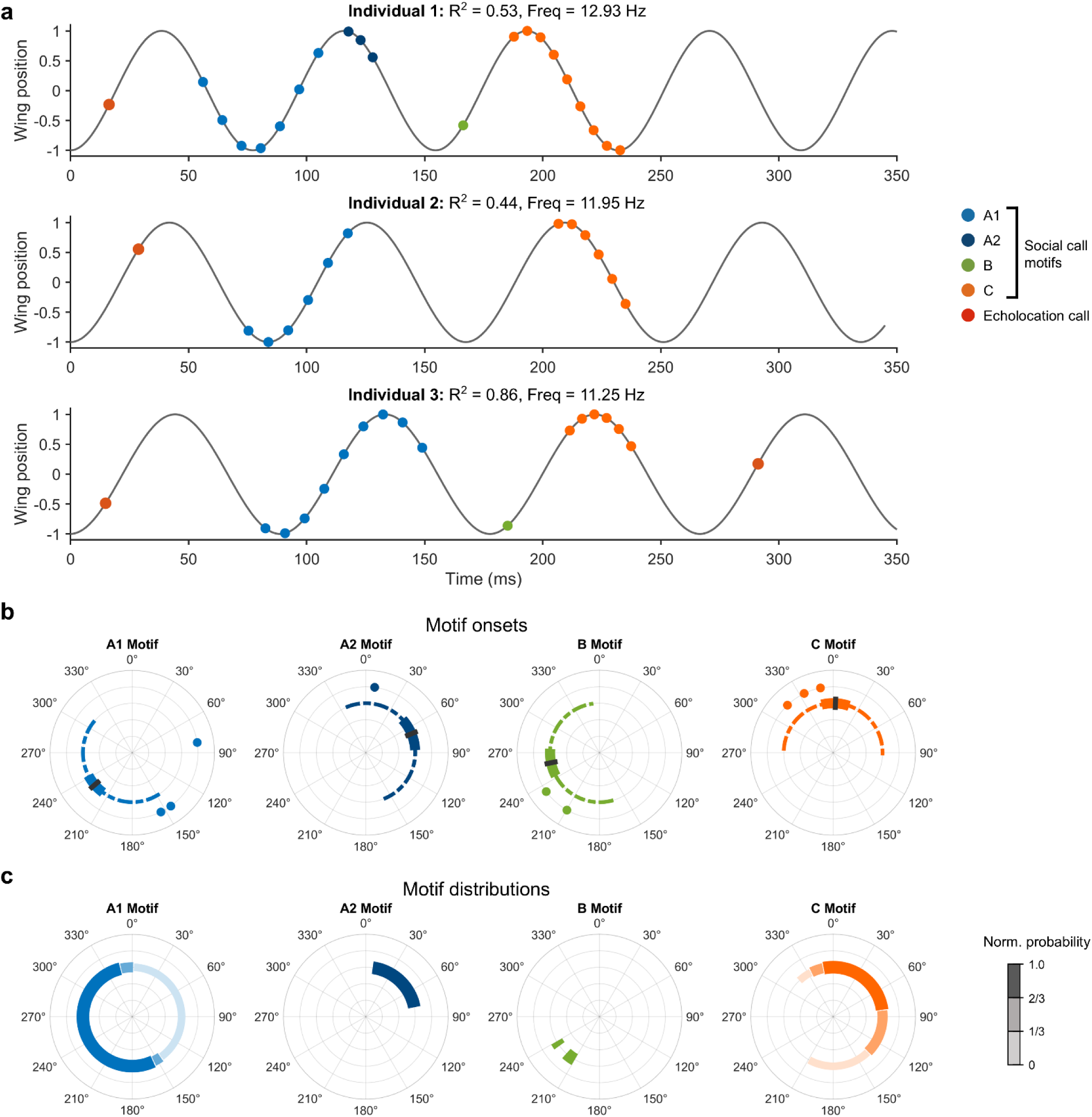
Video-tracked social calls confirm model predictions of stereotyped motif emission. **a** Reconstructed wingbeat cycles of three individuals with various wingbeat frequencies. All recorded acoustic elements of the social call interval are mapped onto each individual’s respective wingbeat cycle. **b** Circular plots showing measured motif onsets of the three individuals from **a** relative to wingbeat phase (colored circles). For each motif, model predictions from Fig. 5b are shown as circular bars for direct comparison (black vertical bars = mean predicted onsets; thick horizontal bars = SEM; dashed horizontal bars = standard deviation). **c** Circular plots showing measured distribution of full motifs relative to wingbeat phase of the three individuals from **a**. For each motif, normalized probability of the respective motif being present at any given wingbeat phase is shown. 0°–180° denotes downstroke, 180°–360° denotes upstroke phase of the wingbeat.

The results of our video tracking confirmed the predictions of our inference model, demonstrating that the complex social call is tightly linked to motor behavior during flight: motif onsets were preferentially emitted at specific phases of the wingbeat cycle, with the SI spanning three consecutive wingbeats. Combined, these results show that vocal-locomotor coupling does not only apply to echolocation in bats, but also acoustic social communication.

## Discussion

In the current study, we investigated how complex social vocalizations are integrated into the locomotor rhythm of flight in wild Nathusius’ pipistrelles (*Pipistrellus nathusii*). We found that the underlying wingbeat period during flight determines the duration of the social call interval, precisely spanning three complete wingbeat cycles. These complex social calls are seamlessly embedded within the regular echolocation sequence, allowing for a transition back to echolocation calls without any temporal disruption. By utilizing a mechanistic inference model and subsequent video tracking, we show that motif onsets are phase-locked to a stereotyped phase of the wingbeat cycle. Together, these results show that bats integrate complex social communication into the wingbeat cycle, and by extension the respiratory rhythm, demonstrating that vocal-motor coupling in bats extends beyond echolocation.

### Vocal-locomotor coupling in bats differs from vocal learner birds

The synchronization of locomotion and respiration is ubiquitous among most tetrapods, likely dating back to the first land-dwelling quadruped tetrapods (Boggs, 2002; Pouw & Fuchs, 2022). In species that vocalize during locomotion, this synchronization extends to vocalizations: the muscles involved in locomotion modulate expiration, which then governs the emission of vocalizations (Butler & Woakes, 1990; Cooper & Goller, 2004; Lancaster et al., 1995; Pouw et al., 2021; Pouw & Fuchs, 2022; Roberts, 1972; Suthers et al., 1972; Teipel & Goller, 2025).

In birds, flight poses unique physiological effects on vocalizations emitted on the move. As birds represent the only other flying vertebrate clade next to bats, they offer a unique comparison when it comes to locomotor-respiratory-vocal coupling during flight (Smotherman et al., 2016). A recent large comparative study in birds showed that species which learn or modify their vocalizations through input of conspecifics, so-called vocal learners, decouple the emission of calls from the wingbeat during flight, whereas the calls of vocal non-learners remained coupled (Berg et al., 2019). This suggests that the emergence of vocal learning and complex vocalizations required vocalizations to become independent from the constraints of locomotion, similar to what has been observed in humans (Pouw & Fuchs, 2022).

Based on their complex social vocalizations, often described as “song” (Jahelková et al., 2008; Smotherman et al., 2016), Nathusius’ pipistrelles have been suggested to be likely candidates for vocal learning (Knörnschild, 2014). Although this hypothesis remains to be tested, the complexity of their social calls, as well as the variable social function of these calls, suggests a decoupling from locomotor patterns, similar to the findings in birds (Berg et al., 2019).

Despite bats being capable of remarkable vocal control (Elemans et al., 2011; Moss et al., 2006; Stidsholt et al., 2021), our findings reveal that this is not the case: instead, we found that complex social vocalizations, consisting of various multi-syllable motifs, are perfectly coupled to the wingbeat during flight. The emission of a single social call always spanned 3 consecutive wingbeats during flight, ensuring a seamless integration of the social call into the regular sequence of echolocation calls. Besides not causing any temporal disruption on the sequence of echolocation calls, this coupling has a second, probably more important consequence: as the social call interval becomes longer, governed by the underlying wingbeat, the onsets of each motif also shift in time. Thus, motif onsets are always emitted around the same particular wingbeat phase, as shown by our model and confirmed by our tracking data. This suggests that while vocal-learner birds relaxed locomotor-vocal coupling in order to gain complexity, bats might have exploited the existing vocal-motor scaffold to build theirs.

### Flight as energetic opportunity

If vocal learner birds decouple their call emission from the wingbeat (Berg et al., 2019), and bats are capable of elaborate vocal control (Elemans et al., 2011; Moss et al., 2006; Stidsholt et al., 2021), then why do we see such strong coupling when it comes to complex social calls during flight in bats?

The most likely explanation is energetic considerations. Recently, Chaverri et al. (2021) showed that the stationary emission of social calls in Spix’s disc-winged bats (*Thyroptera tricolor*) is extremely energetically costly. Emitting social calls more than doubled the average metabolic rate, with vocalizations leading to an estimated instantaneous 26-fold increase in metabolic costs.

Coupling call emission to the wingbeat during flight might mitigate these high costs (Smotherman et al., 2016): the major muscles powering flight, as well as the lateral abdominal wall, contract during the late upstroke phase of the wingbeat (Lancaster et al., 1995). These contractions cause an increase in subglottal pressure, leading to expiration and the emission of vocalizations, thereby lowering energetic effort (Fattu & Suthers, 1981; Lancaster et al., 1995; Suthers et al., 1972). This idea is supported by findings made decades ago for echolocation calls: the stationary emission of echolocation calls in common pipistrelles (*P. pipistrellus*) increased basal metabolic rates by almost 10-fold (Speakman et al., 1989). Importantly, once emitted during flight, echolocation calls did not increase energy expenditure or metabolic rates in comparison to non-vocalizing individuals (Speakman & Racey, 1991; Voigt & Lewanzik, 2012). This shows that flight mitigates the otherwise high costs of echolocation call emission. As a result of this energetic optimization, echolocation call emission across various species is tightly linked to the wingbeat during search flight, usually confined to a narrow time window at the end of the upstroke (Britton et al., 1997; Falk et al., 2015; Koblitz et al., 2010; Moss et al., 2006; Schnitzler, 1971; Stidsholt et al., 2021; Wong & Waters, 2001; Xia et al., 2025). Notably, we found the same coupling in wild Nathusius’ pipistrelles. Echolocation calls were mostly emitted during the latter half of the upstroke, right before the turning point into the downstroke. Conversely, not a single call was emitted during the latter half of the downstroke, where inhalation is supposed to happen (Suthers et al., 1972).

Based on the high costs associated with social call emission during stationary displays (Chaverri et al., 2021), and the tight coupling observed for echolocation calls, we hypothesize that the same mechanism applies to highly complex, song-like vocal displays during flight. Coupling allows bats to produce complex song with little to no additional metabolic cost beyond the cost of flight itself.

### Honest signaling and the "handicap" of the downstroke phase

If the social call is coupled to the wingbeat for energetic reasons, then why do we see emission of the A2 and C motif during the energetically unfavorable downstroke phase? We propose that these motifs might serve as honest signals in Nathusius’ pipistrelles, conveying information about the quality and fitness of a particular male (Zahavi, 1975).

Bat social calls often play an important role in territorial defense (Barlow & Jones, 1997; Götze et al., 2020; Wright et al., 2014) and mate attraction during courtship (Gillam & Fenton, 2016). The social call of *P. nathusii* serves in a dual function, playing a crucial role in intra- and intersexual communication (Jahelková et al., 2008). In order to assess the quality of a potential rival or mate, costly signals can serve as honest signals of the emitter’s fitness (Zahavi, 1975). In bats, such signals can prevent the potentially high costs of physical confrontation with rivals (Bradbury & Vehrencamp, 2011; Zhao et al., 2018) and optimize female mate choice (Behr et al., 2006; Davidson & Wilkinson, 2004). In multiple species of bats, various acoustic and temporal features of social calls were shown to directly predict individual fitness, dominance and aggression in agonistic encounters (Walter & Schnitzler, 2019; Zhao et al., 2018), as well as harem size and reproductive success (Behr et al., 2006; Davidson & Wilkinson, 2004).

The social call motifs A2 and C emitted by *P. nathusii* during the downstroke, and thus at an energetically unfavorable point in the wingbeat, likely represent such an honest signal: individual males that produce longer and more syllable-rich motifs during the downstroke phase of the wingbeat could directly signal their fitness. Longer motifs during the downstroke must either lead to a subsequently shortened inspiratory phase or to postponed exhalation, thus causing higher energetic costs (Fattu & Suthers, 1981; Lancaster et al., 1995; Suthers et al., 1972).

In line with this hypothesis, the C motif typically exhibits narrow within-individual variation, while showing large variation between individuals, indicative of an individual fitness signal (Jahelková et al., 2008). Moreover, the number of C syllables emitted by any given individual increases when a foreign male of the same species enters its territory (Jahelková et al., 2008).

The A2 motif on the other hand is emitted directly after the A1 motif without any major pause. However, A2 emission varies a lot between individuals: in some, its power is so weak, that its either impossible to record or might not even be emitted at all (Jahelková et al., 2008). Accordingly, we only recorded it in 68% of all social calls. This suggests that only the fittest males might be able to emit an A2 motif during the downstroke phase. Moreover, a longer A1 motif would subsequently shift A2 emission further towards the downstroke phase. Matching this idea, A1 duration and syllable number reportedly increases during agonistic encounters (Jahelková et al., 2008).

### Evolution of social calls and the emergence of call rhythms

The coupling of social call emission and wingbeat that we observed in *P. nathusii* should not be viewed as a simple physiological constraint. Instead, this coupling might have created the energetic conditions that allowed complex vocal signals to emerge in the first place. Smotherman et al. (2016) suggested that higher efficiency through vocal-locomotor integration during flight could explain why elaborate song remains rare in most mammals. In this sense, coupling during flight might represent an evolutionary opportunity rather than a limitation.

Several pipistrelles closely related to *P. nathusii* share the same basic social call element, namely the A motif (Agnarsson et al., 2011; Dool & Puechmaille, 2025; Pfalzer & Kusch, 2003). Most of them also emit this motif during flight, where it fulfills similar social functions (Smotherman et al., 2016). This suggests that some precursor of the A motif most likely existed in the ancestor of all western pipistrelles (Dool & Puechmaille, 2025; Zhukova et al., 2023), likely as a flight-emitted vocalization synchronized to the wingbeat cycle for energetic reasons. In *P. nathusii*, this ancestral motif subsequently evolved into a more elaborate vocal display. However, the display remained tightly embedded within the locomotor-respiratory rhythm of flight, as shown by our findings. This suggests that *P. nathusii* did not evolve complexity by escaping the locomotor scaffold, but rather by exploiting it (Pouw et al., 2021; Smotherman et al., 2016). It remains an open question why P. nathusii stands out within its genus as the only species to have expanded upon this ancestral motif, evolving multi-motif, song-like signals.

This evolutionary origin in flight also helps explain another striking observation: Jahelková et al. (2008) found no major differences in spectral or temporal characteristics when comparing social calls of *P. nathusii* emitted during flight to those produced during stationary displays. Thus, the rhythmic organization of the call likely evolved under flight and remained conserved even when its use shifted to also include sedentary displays.

More broadly, this fits into a wider evolutionary scenario for bat vocal communication. Flight likely preceded the evolution of high-frequency echolocation (Liu et al., 2022; Simmons & Geisler, 1998; Speakman, 2001; Teeling et al., 2012; Wang et al., 2017). As echolocation evolved as a means of orientation during nocturnal flight, energetic constraints tightly coupled call emission to wingbeat and respiratory rhythms. As these echolocation calls differentiated into more complex social vocalizations, the underlying rhythms could have remained anchored to the same locomotor-respiratory scaffold, similar to what we observed in *P. nathusii* (Burchardt et al., 2019; Pouw et al., 2021; Smotherman et al., 2016). This persistence of rhythmic coupling even outside of flight also finds support in other bat species. Greater sac-winged bats (*Saccopteryx bilineata*) emit various social calls solely during stationary displays, where they follow an isochronous rhythm, similar to respiratory and wingbeat rhythms (Burchardt et al., 2019). Isochronous rhythms in sedentary displays could therefore represent vestigial traits of an ancestral flight-coupled state.

The persistence of these rhythmic patterns, even in the absence of flight, might also be rooted in the underlying neural architecture of the brain. Across mammals, central pattern generators coordinate locomotor and respiratory rhythms (Bass, 2014; Giraudin et al., 2012; Le Gal et al., 2020), and vocal premotor neurons are tightly gated by respiratory circuits, ensuring that phonation remains locked to expiration (Park et al., 2024). This organization implies that vocal rhythms are inherently embedded within respiratory and locomotor dynamics. Consequently, if flight entrains respiration, both echolocation and social calls should remain temporally anchored to this locomotor-respiratory framework. In bats, echolocation and social communication calls seem to be controlled by partly overlapping brainstem and midbrain circuits whose activity patterns are flexibly reconfigured according to context (for review, s. Babl et al., 2025). However, how the brain supports and structures these two vocal modes remains a major open question in the field (Babl et al., 2025).

### Conclusion

In conclusion, our results show that complex social calls in bats are coupled to the mechanics of flight, and thus likely also to respiratory patterns. Unlike vocal-learning birds, bats do not show vocal-locomotor decoupling during complex vocalizations. Instead of constraining vocalizations, flight might have represented an opportunity for bats to evolve complex social signaling in the first place by reducing the energetic cost of vocal signaling. Thus, biomechanical integration might have actively shaped the evolution of one of the most elaborate vocal communication systems in the animal kingdom.

## Acknowledgments

We thank Sabine Horlacher and the team of the Federsee-museum in Bad Buchau, Germany, for access to their property over the years. We also thank Dr. Diana Schöppler for her support, Prof. Dr. Hans-Ulrich Schnitzler for providing recording equipment and Prof. Dr. Marc Schmidt for comments on the topic.

This work was supported by the Germany scholarship program of the University of Tübingen in collaboration with the Wolfgang Rosenstiel Foundation and the Gips-Schüle-Foundation, and by a scholarship of the Reinhold-und-Maria-Teufel-Foundation to SB.

## Author contributions

SB, LR and AD designed the study. SB and LR collected and analyzed the data. SB performed the coding and statistical analyses, created figures, and wrote the original manuscript. LR and AD reviewed and edited the manuscript.

## Data and code availability

All data and analyses code will be made available to the public after publication.

