## Supplemental Information for "Complex Social Vocalizations are Integrated into the Bat Wingbeat Cycle during Flight"

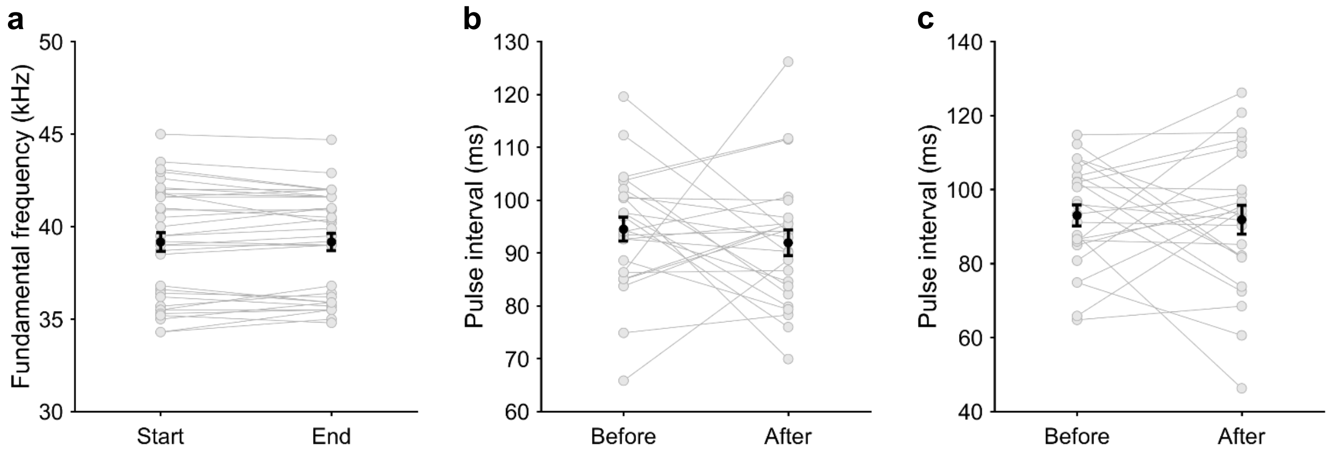


**Figure S1. Echolocation call parameters stay consistent before and after social call emission. a** Terminal frequency of echolocation calls (ECs) before and after the social call. **b** Mean EC pulse interval before and after the social call. **c** EC pulse interval immediately before and after the social call.

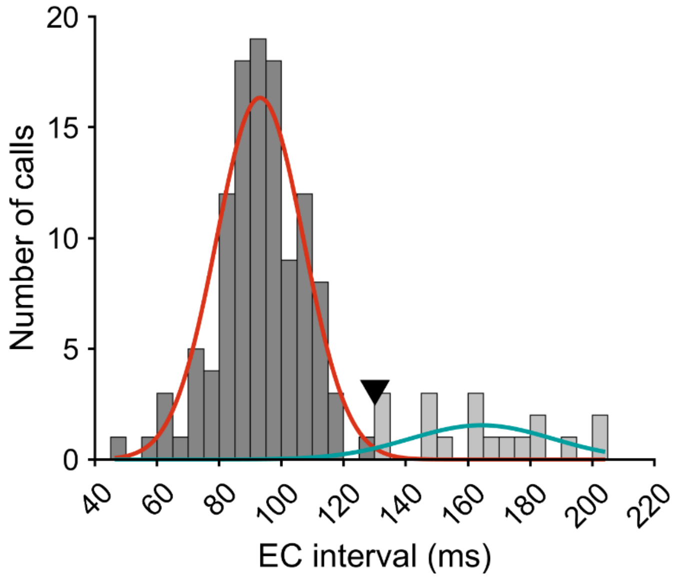


**Figure S2. Bimodal distribution of echolocation pulse intervals.** As echolocation call emission is occasionally omitted for a single wingbeat, pulse intervals exhibit a bimodal distribution. The two best fitting gaussian distributions for the data were estimated via a Gaussian Mixture Model, illustrated in red and turquoise. Crossover point of the two distributions is shown as black triangle.

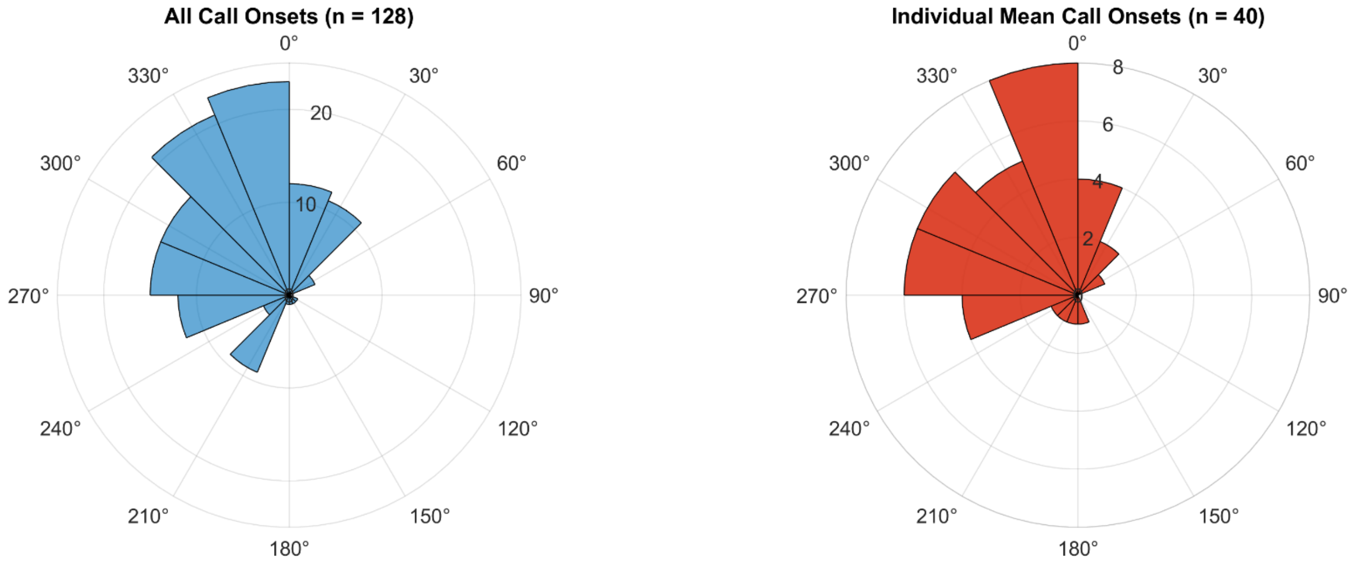


**Figure S3. Polar histograms of reconstructed wingbeat phases during echolocation call emission for all calls and population data.** Left shows polar histogram of wingbeat phases for all measured calls, right for population data, where each sample represents the mean of an individual recording. 0°–180° denotes downstroke, 180°–360° denotes upstroke phase of the wingbeat.
